# On-body Measure of Reaction Time Correlates with Intoxication Level

**DOI:** 10.1101/2025.04.17.649358

**Authors:** Megan Blackwell, Henry Valk, Josh Eaton, Sam Kovaly, Sam Karnes, Tue Vu, Kate Stevenson, Mike Kowalczyk, William Gormley

**Affiliations:** Pison; Massachusetts General Brigham

## Abstract

Excessive alcohol use has profound effects on individual health and healthcare systems worldwide. Despite this, there is currently no system or device that can detect robustly the physiologic and functional effects of alcohol-based impairment in real-world conditions. A practical, on-body, device capable of rapidly and accurately determining the functional capacity of an individual to drive, before they can start the ignition of the automobile, is required. The goal of this pilot study was to evaluate the effect of acute alcohol intoxication on premotor time (PMT) and reaction time (RT), both highly sensitive of individual cognition, as evaluated by the Pison Technology wrist-worn wearable. Nineteen participants were included in the study, 14 subjects who consumed alcohol sufficient to raise blood alcohol concentration (BAC) to 0.12% within a 30-minute period and 5 controls who did not consume alcohol. Changes in reaction time data were correlated with blood alcohol levels as measured by breathalyzer testing, identifying a statistically significant difference between those participants under the legal limit and those over the legal limit. Both group and individual analyses confirmed that as the BAC increased in subjects, the PMT also increased. The PMT also decreased as the BAC returned to levels under the legal intoxication threshold (0.08%). A significant effect of BAC levels on changes in PMT at the p < 0.05 level for two conditions. This study demonstrated the first use case of an on-body, neuro-physiological sensor capable of detecting sensitive changes of reaction time, in real time, that serves as an easy-to-measure proxy for blood alcohol content and impairment.

## Introduction

According to the Center for Disease Control, excessive alcohol use resulted in over 140,000 deaths and 3.6 million years of potential life lost each year in the United States from 2015 to 2019, and over 178,000 deaths and 4.3 million years of potential liwith younger populations bearing a disproportionate portion of this risk.^1^ Worldwide, 13.5% of total deaths in the 20-39 age group were attributed to alcohol use.^2,3^ In 2020 alone, there were 11,654 fatalities in traffic crashes in which at least one driver was alcohol-impaired, with a blood-alcohol concentration (BAC) exceeding 0.08 g/dL.^4^ This seemingly preventable loss of life motivated the 2021 Infrastructure Investment and Jobs Act to require inclusion in all new passenger cars a technology that can “passively monitor the performance of a driver … to accurately identify whether that driver may be impaired” and “prevent or limit motor vehicle operation if an impairment is detected.”^5^ To achieve the vision of eliminating drunk driving, it requires the development of technologies that are easy to use, self-effacing, and provide user-specific data in a timely fashion. Despite the rapid advancement and proliferation of sensor technology, there is currently no system or device that unobtrusively and inconspicuously assesses impairment created by alcohol consumption to 1) enable users to refrain from consuming more alcohol when impairment is detected and 2) give individuals quantitative data on their ability (or lack thereof) to manage equipment, operate machinery, and drive.

Although two emerging technologies were identified by the National Highway Traffic Safety Administration and the Driver Alcohol Detection System for Safety as promising candidates for ignition interlock devices, each presents challenges in the robust detection of intoxication.^6^ TruTouch, a technology that uses diffuse reflectance spectroscopy at the finger, suffers from weak ethanol absorption and confounding absorptions from other skin tissue components.^7^ Measurements from Autoliv, a technology that uses non-dispersive infrared gas sensors on ambient air, have a large variability due to challenges in sample collection and complex calibration.^8^ It also seems that an open car window would compromise Autoliv’s sensitivity. These passive devices do not require active participation by drivers, but each presents detection challenges. This suggests that existing wearable alcohol sensors cannot be used as an effective ignition interlock device. **Consequently, there remains an unmet need for an easy-to-use yet robust sensor to detect alcohol impairment.**

A direct and more reliable way to measure impairment associated with increased blood alcohol levels is to measure reaction time rather than measuring alcohol itself. Reaction time is crucial in preventing motor vehicle accidents.^21,22^ Studies have investigated driver reaction time to avoid a collision and measured a mean time of 2.2 seconds required to apply brakes and 1.6 seconds to steer away.^11^ Several studies have measured the relationship between blood alcohol concentration (BAC) and reaction time, linking increasing alcohol intake with slower stimulus-response behavior.^10, 12–14^ A BAC of 0.06% increased brake reaction time by 20%.^15^ Reaction time alterations are widely understood to be altered early and reliably in cases of alcohol intoxication.

Pison has recently developed a wearable device that can detect increases in reaction time, which accompany any intoxication, specifically ones caused by alcohol. Intoxication assessment can be accomplished by a patient-initiated reaction time test and the results compared to a baseline value. One envisioned end use of the Pison wearable device provides users with direct information to promote healthy choices concerning alcohol use and increase public safety. Another end use advances the device as an alcohol or fatigue interlock for motor vehicles. Pison aims to commercialize the first ignition-interlock device that infers impairment from increased reaction time measurements acquired on its wrist-worn sensor. Another envisioned use for this technology is by transportation organizations that wish to assess drivers or operators for readiness to increase the safety of operations.

This study aims to quantify the accuracy of reaction time-based models to predict blood alcohol concentration in real time. The study is a prospective, non-randomized, single-arm intervention designed to collect laboratory, physiologic, and psychometric subject data during progressive administration of weight- and gender-based oral ethanol administration. In this pilot study, we collected physiologic and behavioral biomarkers using the Pison device to investigate the relationship between blood alcohol levels and cognitive and physical impairment as measured by reaction time and electroencephalography (EEG). Results from the EEG study will be reported in a separate paper.

The Pison wearable collects multi-channel surface electromyography (EMG) measurements from the peripheral nervous system. The Pison wearable used in this study is shown in Figure 1 and includes five channels of biosignal data measured from differential pairs of electrodes on the surface of the skin. Filtering and preprocessing of sensor data are performed on device, resulting in high-quality EMG signals. The Pison wearable is paired to a compatible mobile phone to which sensor readings are shared via Bluetooth.

**Figure 1:**
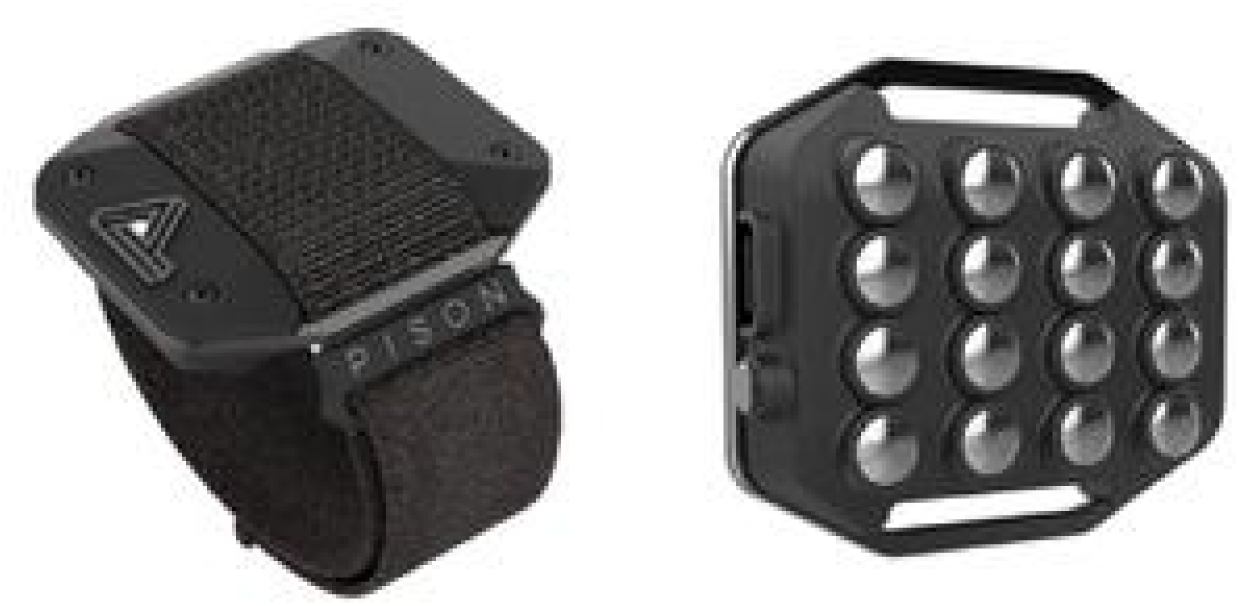
Pison Wearable Device. Pison’s sensor is worn on the wrist and uses stainless steel electrodes to measure biopotential activity on the top of the wrist.

**Figure 2:**
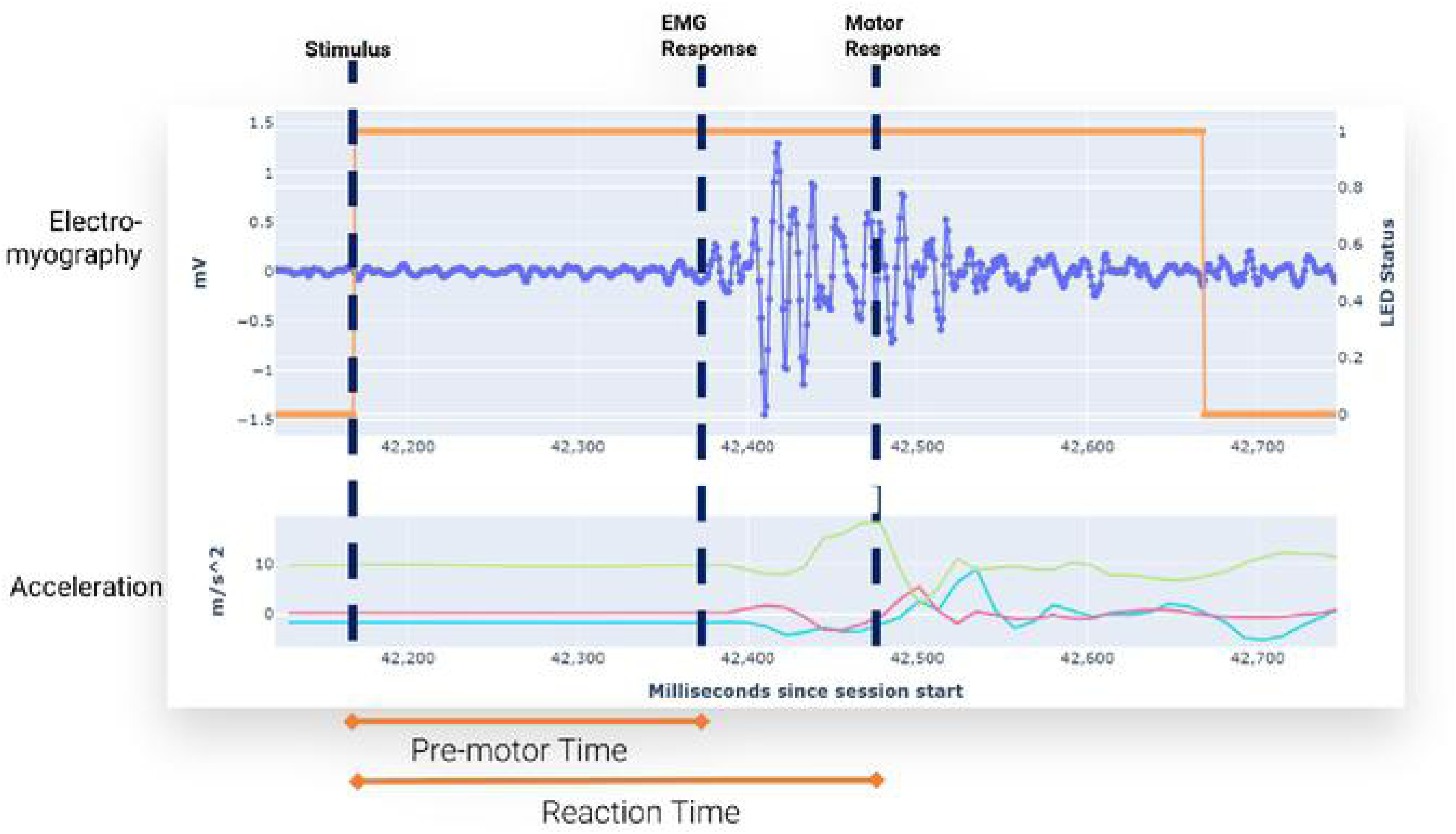
The Pison wearable uses surface electromyography (EMG) and accelerometry data to measure premotor time (PMT) in response to a visual stimulus presented on the device. Premotor time describes the time elapsed between the visual stimulus and the neuromuscular response as measured by an increase in the EMG signal. Response onset is determined by Pison’s proprietary algorithm. Reaction time is determined by measuring the onset of movement as measured by the accelerometer signal. Electromechanical delay describes the difference between the pre-motor time and the reaction time.

The Pison wearable uses EMG and accelerometry data to measure premotor time (PMT) and reaction time (RT) in response to a visual stimulus presented on the device. Premotor time describes the time elapsed between the visual stimulus and the neuromuscular response as measured by an increase in the EMG signal. Reaction time is determined by measuring the onset of movement as measured by the accelerometer signal. Electromechanical delay is defined as the time between EMG onset and the onset of force or motion and is estimated by the difference between the measured premotor and the reaction times.^16^ The Pison reaction time assessment is entirely self-contained and does not rely on touchscreens or peripheral devices that can suffer from latencies due to operating system delays.^17^

A custom software application (Figure 3) running on a mobile device, such as a cell phone or tablet, displays premotor time and reaction time results as well as comparisons with baseline performance. A user may opt-in to share data with others who have proper permissions.

**Figure 3:**
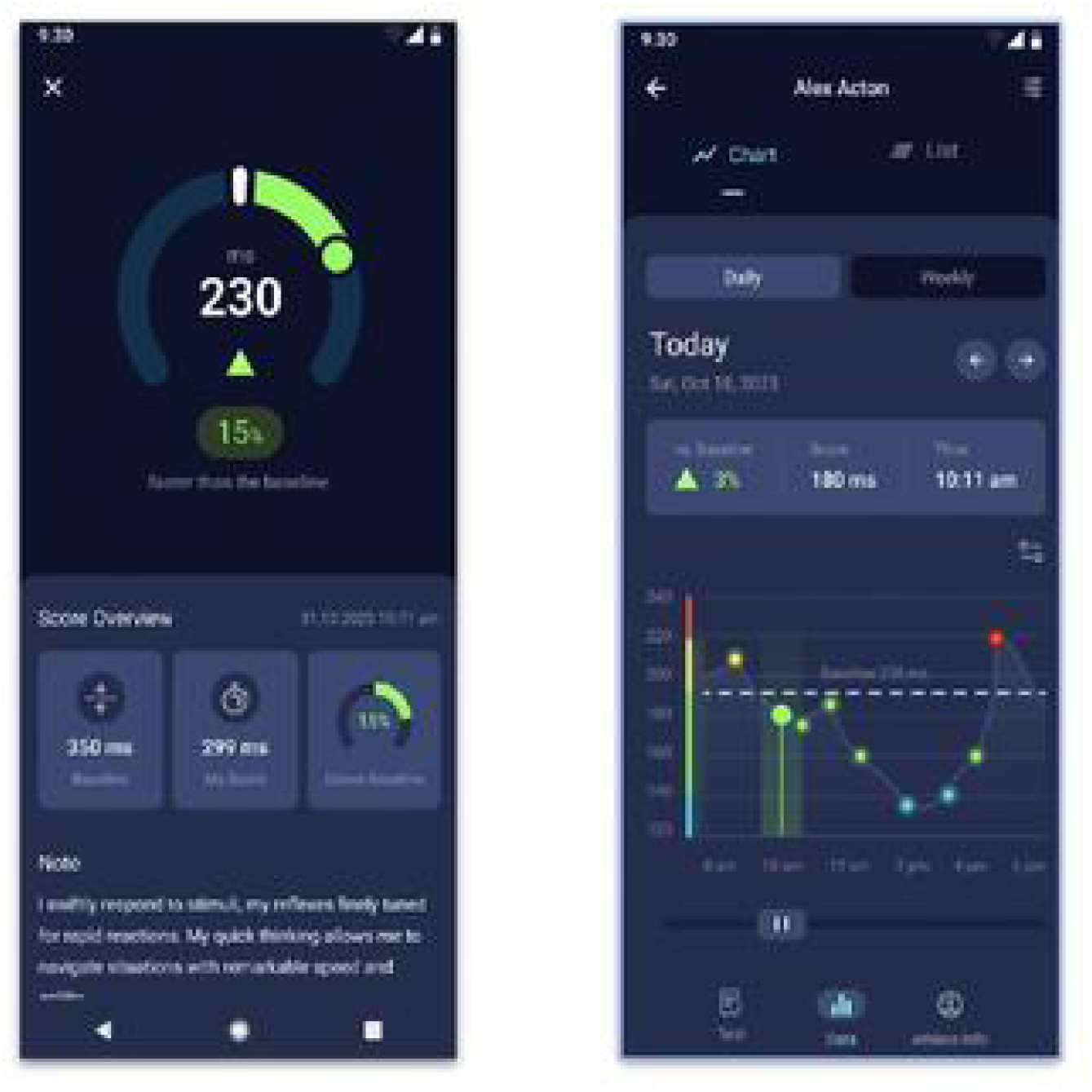
**Pison Mobile Application. Pison’s mobile application quantifies readiness by comparing reaction-time measurements to a baseline data set.**

This study demonstrates the use of reaction time to measure impairment related to levels of blood alcohol exceeding the legal limit. The neurophysiologic data measured with the Pison wearable provides a robust predictive capacity to determine, at a population level, in a statistically significant manner, the difference between subjects below and above the legal BAC. Additionally, at an individual level, a very robust mirroring of reaction time to blood alcohol level was shown.

## Materials and Methods

### Participants & Protocol

The Pison study was approved by the Western-Copernicus Group (WCG) Institutional Review Board (IRB). A total of 14 test subjects (6 males, 8 females) and five controls (3 males, 2 females) ranging in age from 21-36 years were recruited between September 26 and October 3, 2023; there were 8 males and 10 female participants who completed testing. Subject enrollment occurred after written informed consent was obtained. Those excluded from the study were students, active-duty military personnel, prisoners, cognitively impaired persons, women who were pregnant, and individuals who had backgrounds of alcohol or substance abuse. All subjects were screened for alcohol dependence through the Alcohol Dependence Scale during intake and excluded if scoring 8 or above. Women who consented to participate were provided a pregnancy test to be taken on-site. Subjects were asked to step on a scale to ascertain their body weight in pounds to support the calculation of a standardized and personalized alcohol dosage, shown in Table 1. The intent of the dosage was to produce a BAC over 0.10%, but ideally no greater than 0.12%, after 30 minutes from the time of consumption.

**Table 1.** Blood Alcohol Concentration Estimations for (a) men and (b) women. These published estimations were used to guide the number of servings each participant in the study should have in order to achieve the desired blood alcohol level. One drink was equivalent to a 1.25 oz serving of an 80 Proof liquor, or a 12 oz serving of beer, or a 5 oz serving of wine. After each 40 min period of drinking, 0.01 gm% was subtracted from the estimated blood alcohol level.^18^

| Body Weight (lb) |  |  |  |  |  |  |  |  | Body Weight (lb) |  |  |  |  |  |  |  |  |
| --- | --- | --- | --- | --- | --- | --- | --- | --- | --- | --- | --- | --- | --- | --- | --- | --- | --- |
| Drinks | 100 | 120 | 140 | 160 | 180 | 200 | 220 | 240 | Drinks | 100 | 120 | 140 | 160 | 180 | 200 | 220 | 240 |
| 1 | 0.04 | 0.03 | 0.03 | 0.02 | 0.02 | 0.02 | 0.02 | 0.02 | 1 | 0.05 | 0.04 | 0.03 | 0.03 | 0.03 | 0.02 | 0.02 | 0.02 |
| 2 | 0.08 | 0.06 | 0.05 | 0.05 | 0.03 | 0.03 | 0.03 | 0.03 | 2 | 0.09 | 0.08 | 0.07 | 0.06 | 0.05 | 0.05 | 0.04 | 0.04 |
| 3 | 0.11 | 0.09 | 0.08 | 0.07 | 0.06 | 0.06 | 0.05 | 0.05 | 3 | 0.14 | 0.11 | 0.10 | 0.09 | 0.08 | 0.07 | 0.06 | 0.06 |
| 4 | 0.15 | 0.12 | 0.11 | 0.09 | 0.08 | 0.08 | 0.07 | 0.06 | 4 | 0.18 | 0.15 | 0.13 | 0.11 | 0.10 | 0.09 | 0.08 | 0.08 |
| 5 | 0.19 | 0.16 | 0.13 | 0.12 | 0.11 | 0.09 | 0.09 | 0.08 | 5 | 0.23 | 0.19 | 0.16 | 0.14 | 0.13 | 0.11 | 0.10 | 0.09 |
| 6 | 0.23 | 0.19 | 0.16 | 0.14 | 0.13 | 0.11 | 0.10 | 0.09 | 6 | 0.27 | 0.23 | 0.19 | 0.17 | 0.15 | 0.14 | 0.12 | 0.11 |
| 7 | 0.26 | 0.22 | 0.19 | 0.16 | 0.15 | 0.13 | 0.12 | 0.11 | 7 | 0.32 | 0.27 | 0.23 | 0.20 | 0.18 | 0.16 | 0.14 | 0.13 |
| 8 | 0.30 | 0.25 | 0.21 | 0.19 | 0.17 | 0.15 | 0.14 | 0.13 | 8 | 0.36 | 0.30 | 0.26 | 0.23 | 0.20 | 0.18 | 0.17 | 0.15 |
| 9 | 0.38 | 0.31 | 0.27 | 0.23 | 0.21 | 0.19 | 0.17 | 0.16 | 9 | 0.41 | 0.34 | 0.29 | 0.26 | 0.23 | 0.20 | 0.19 | 0.17 |

Two days before the test, eight subjects who had never used the Pison wearable were familiarized with the device and reaction time tests. Baseline data were recorded. The remaining 11 subjects had previously collected baseline data. Subjects who were unable to retrieve a device prior to the day of intervention were asked to take baseline measurements two days after and have been noted in the data.

Subjects were instructed to complete an active RT test or “Salus Protocol” in the morning, afternoon, and evening leading up to the intervention to account for training effects on RT. The “Salus Protocol” includes passive EMG measurements of reaction time, recorded while participants positioned their arm in a “watch-check” position. During each active RT measurement, subjects were prompted by a 500 ms duration white LED light emitted by the wearable and asked to perform the specific action of opening their hand in response to the light as fast as possible. The active test lasted three minutes with a randomized inter-stimulus interval between 1 and 4 seconds.

On the day of the experiment, intake questionnaires were completed and signed in private. All physiological data measures were labeled with the subject’s unique identification number, along with forms regarding the behavioral, physiological, and psychomotor data collected. Participants were instructed to not consume food 8 hours prior to their session in order to expedite the effects of alcohol. Alcohol was administered to participants but not to controls after obtaining baseline measurements for all tasks, in which 100% of the determined dose was consumed over the course of ten minutes. After dosage, a cyclic process of taking breathalyzer (BACtrack S80), EEG (32-channel Enobio, Neuroelectrics), and RT measurements were continued for the remainder of the experiment in alternating blocks performed every 15 minutes, as shown in Figure 4. In the first block, passive EMG, breathalyzer, and RT tests were administered. In the second block, an EEG task was also administered. This resulted in six passive, active, and breathalyzer measurements and three EEG measurements in the 90-minute session. Thirty minutes after the initial dosage, if the participants’ BAC had not reached 0.10% or greater, additional alcohol was administered based on the standardized reference table (Table. 1), taking into account the participant’s current BAC, desired BAC, and body weight.

**Figure 4:**
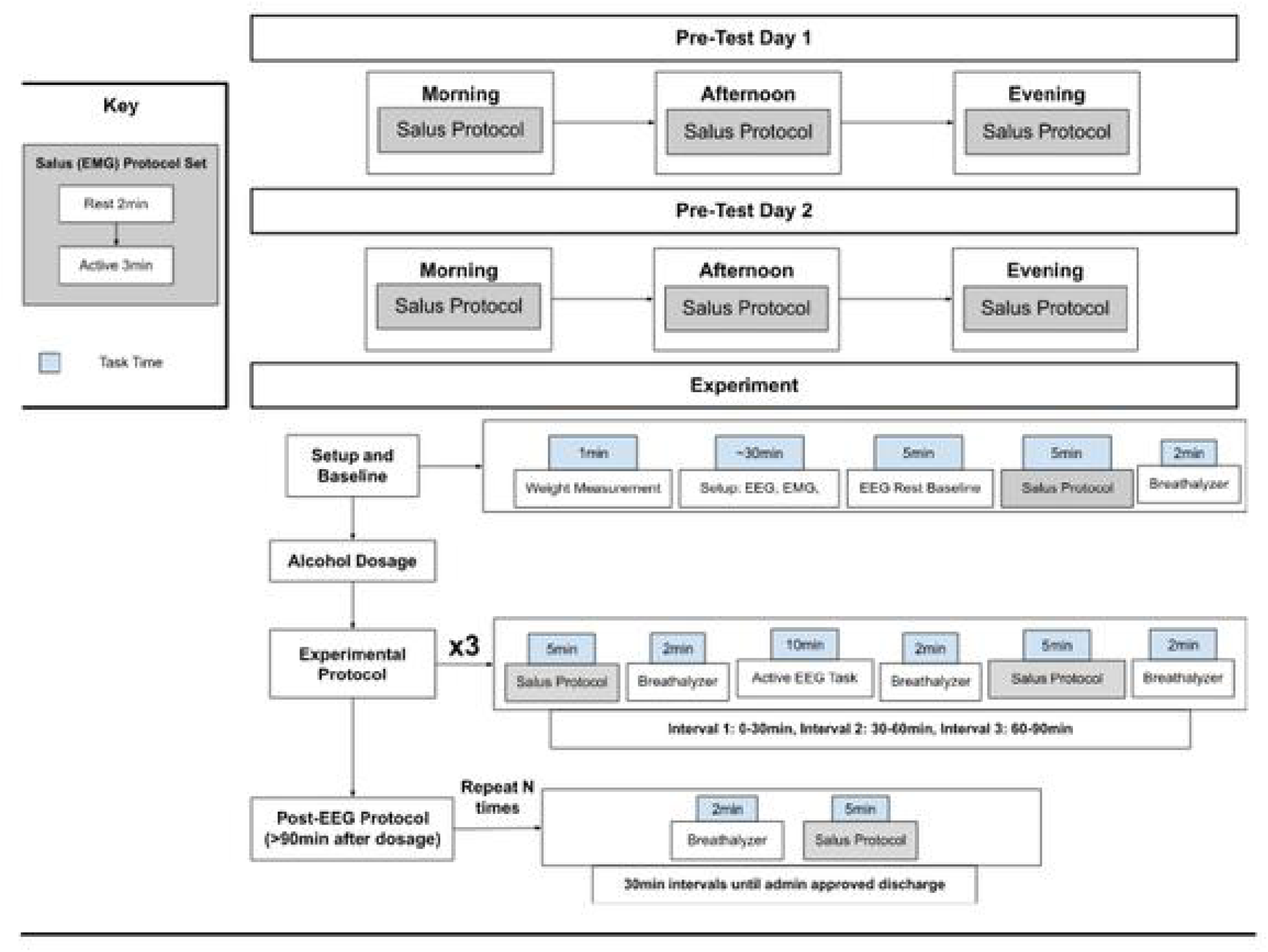
Experimental protocol of BAC study.

The BACTrack breathalyzer is shown in Figure 5A and has a removable mouthpiece that was cleaned or replaced before each participant. The 32-channel Enobio EEG headset is shown in Figure 5B and was used to capture brain activity. Lab Streaming Layer was used simultaneously with the headset’s software to synchronize and record the EEG measurements.

**Figure 5A:**
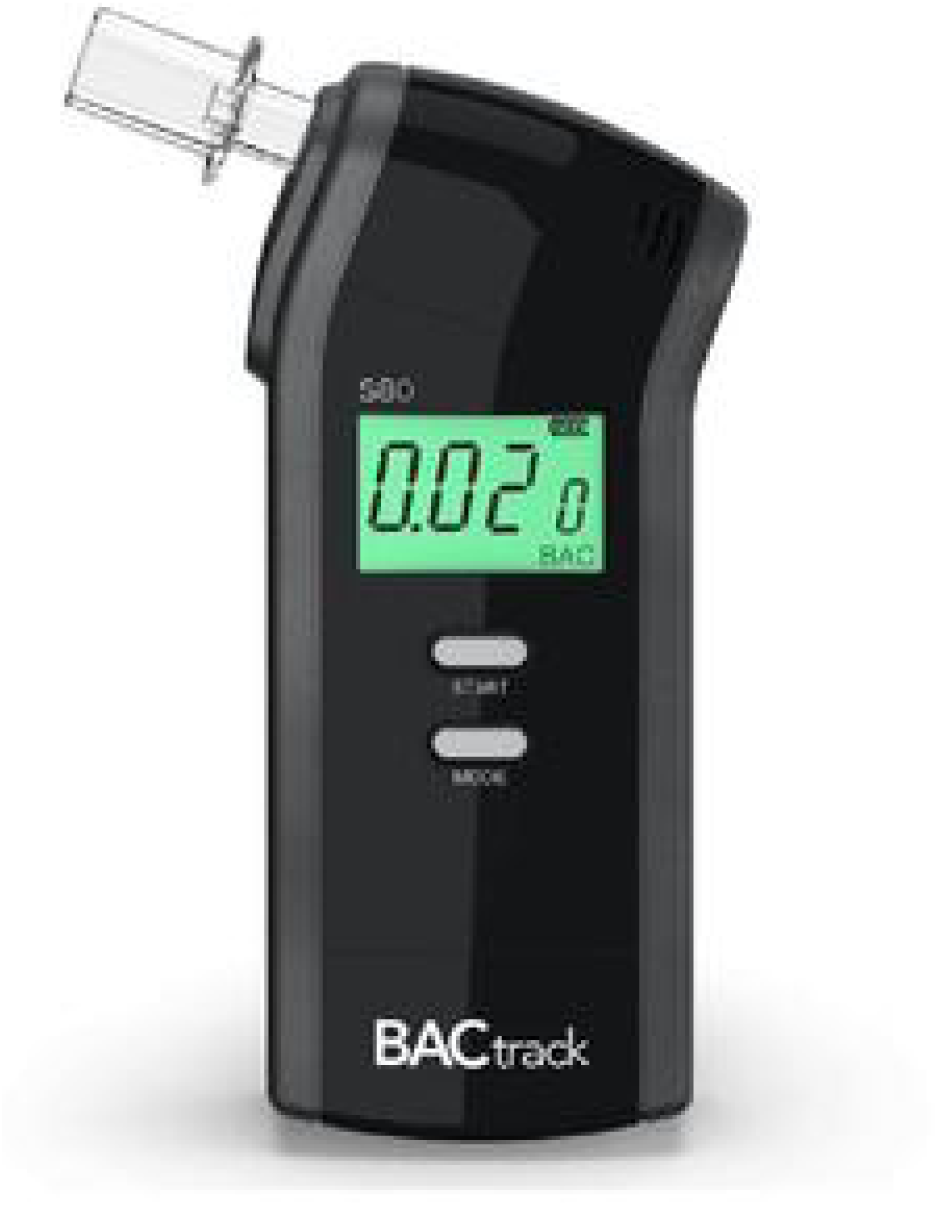
BACtrack S80 (left). BACtrack is the first FDA-approved consumer breathalyzer and has been used for over 20 years in various industries including law enforcement, hospitals, and the military.

**Figure 5B:**
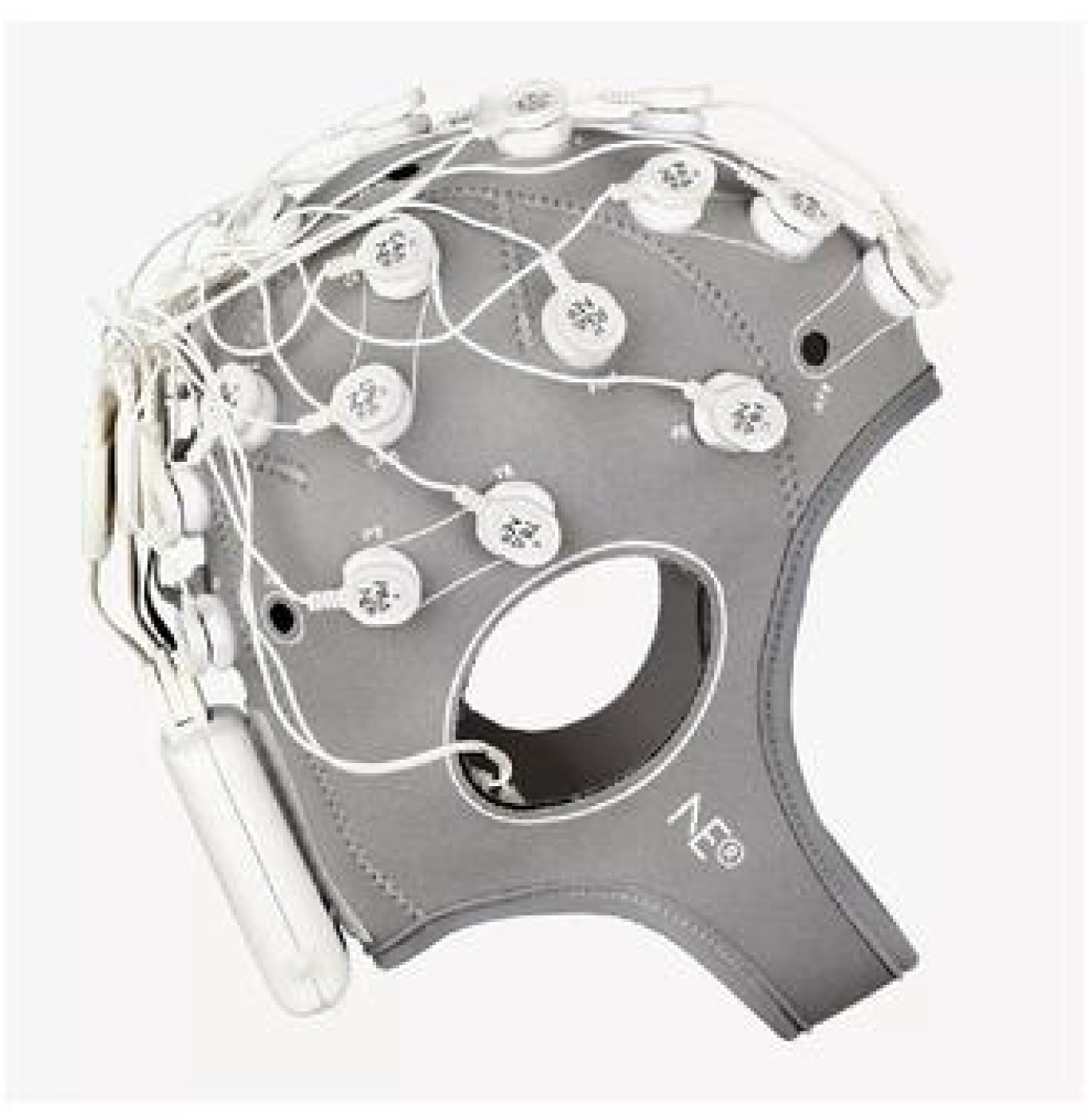
Enobio 32 headset (right). The Enobio 32 is a high-density wireless medical-grade EEG headset from Neuroeletrics. The accompanying software supports LabStreamingLayer.

## Results

### Reaction Time and Blood Alcohol

A one-way between-subjects analysis of variance (ANOVA) was conducted to compare the effect of BAC on changes in reaction time, noted as “Delta RT,” for low (BAC <= 0.05%), medium (0.05% < BAC <= 0.08%), and high (BAC > 0.08%) levels. Changes in RT for the control group were also included in the comparison. BAC levels were found to have a significant effect on Delta RT at the p < 0.05 level for the four conditions [F(3, 3545) = 117, p = 2 e-16], as illustrated in Figure 6. Post hoc comparisons using the Tukey honestly significant difference (HSD) test indicated that the mean score for the low BAC level (M = 67.5, SE = 1.24) was significantly different than the control (M = 72.6, SE = 1.38) with a p-value of 0.03. The mean score for the medium BAC level (M = 104.4, SE = 2.67) was significantly different than the control with a p-value less than 0.05, and the high BAC level (M = 112.7, SE = 2.68) was significantly different than the control with a p-value less than 0.05. Additionally, the medium BAC level was significantly different than the low BAC level and the high BAC level was significantly different than the low BAC level with p-values of less than 0.05. There was not a significant difference between the high and medium BAC levels (p = 0.13). This analysis suggests that elevated BAC levels will increase Delta RT.

**Figure 6:**
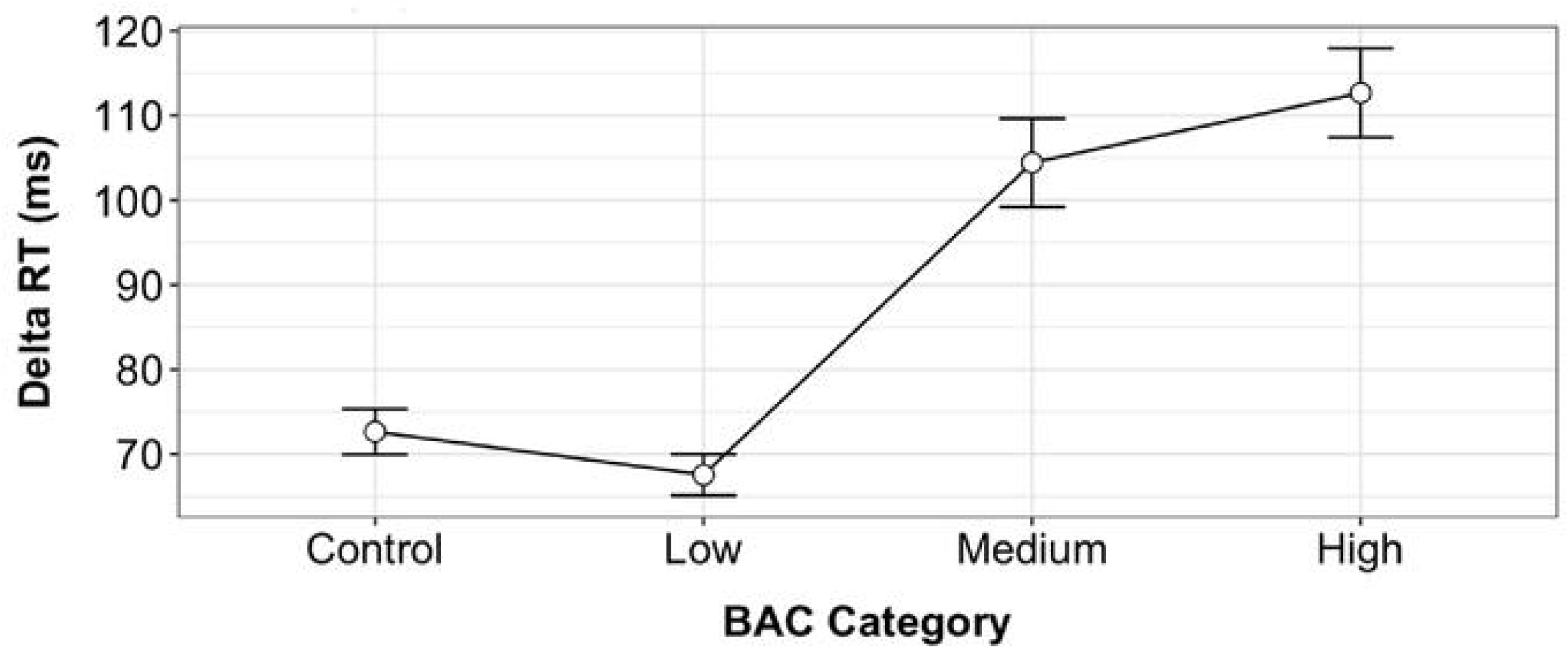
The effect of BAC level on RT. A one-way between-subjects analysis of variance (ANOVA) illustrating the effect of blood alcohol concentration (BAC) on changes in reaction time (RT) for low (BAC <= 0.05%), medium (0.05% < BAC <= 0.08%), and high (BAC > 0.08%) levels. Pairwise comparisons show statistically significant differences between each factor at the p < 0.05 level for the four conditions [*F*(3, 3545)= 117, *p*= 2e-16].

A simple linear regression analysis was conducted on participant data to evaluate if BAC levels could predict Delta RT. A significant regression was found (*F*(1, 3547) = 323.3, *p* = 2.2e-16). The *R*^2^ was 0.084, indicating that the BAC level explained approximately 8.4% of the variance in Delta RT. Figure 7 illustrates the results of this linear model.

**Figure 7:**
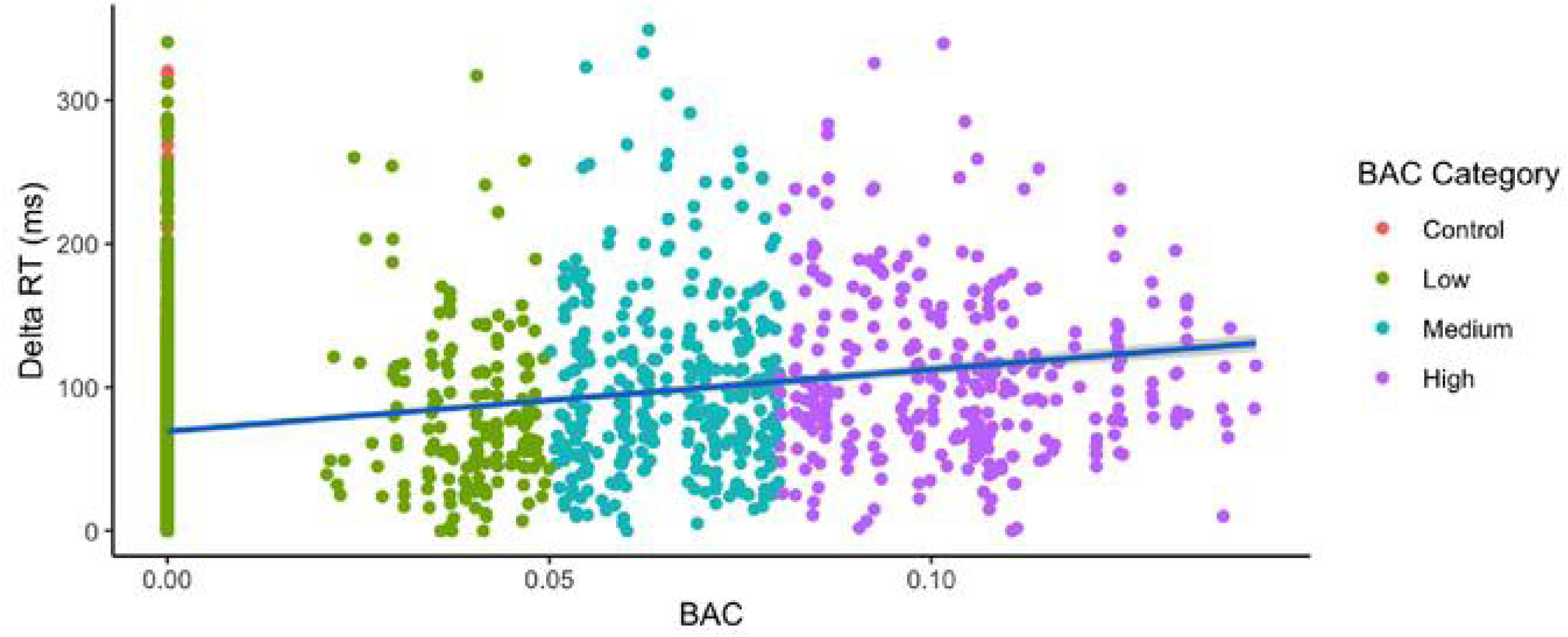
Linear regression analysis demonstrates the ability of blood alcohol content to predict changes in reaction time. Subject data was pooled for all participants and is plotted in red for control subjects, green for low levels (BAC <= 0.05%), teal for medium levels (0.05% < BAC <= 0.08%), and purple for high levels (BAC > 0.08%). A simple linear regression analysis was conducted to evaluate the extent to which BAC levels could predict the change in reaction time, denoted as “Delta RT.” A significant regression was found (*F*(1, 3547) = 323.3, *p* = 2.2e-16). The *R*^2^ was 0.084, indicating that the BAC level explained approximately 8.4% of the variance in delta RT.

A one-way between-subjects ANOVA was conducted to compare the effect of BAC on Delta RT for BAC values under the legal limit (BAC < 0.08) and over the legal limit (BAC >= 0.08). There was a significant effect of BAC levels on delta RT at the p < 0.05 level for the two conditions [F(1, 3547) = 185.3, p = 2e-16]. Post hoc comparisons using the Tukey HSD test indicated that the mean score for the under-legal limit BAC level (M = 73.5, SE = 0.89) was significantly different than the over-legal limit BAC level (M = 112.4, SE = 2.71) with a p=0.0001. These results suggest that increases in BAC levels will significantly increase Delta RT. Figure 8 illustrates these results.

**Figure 8:**
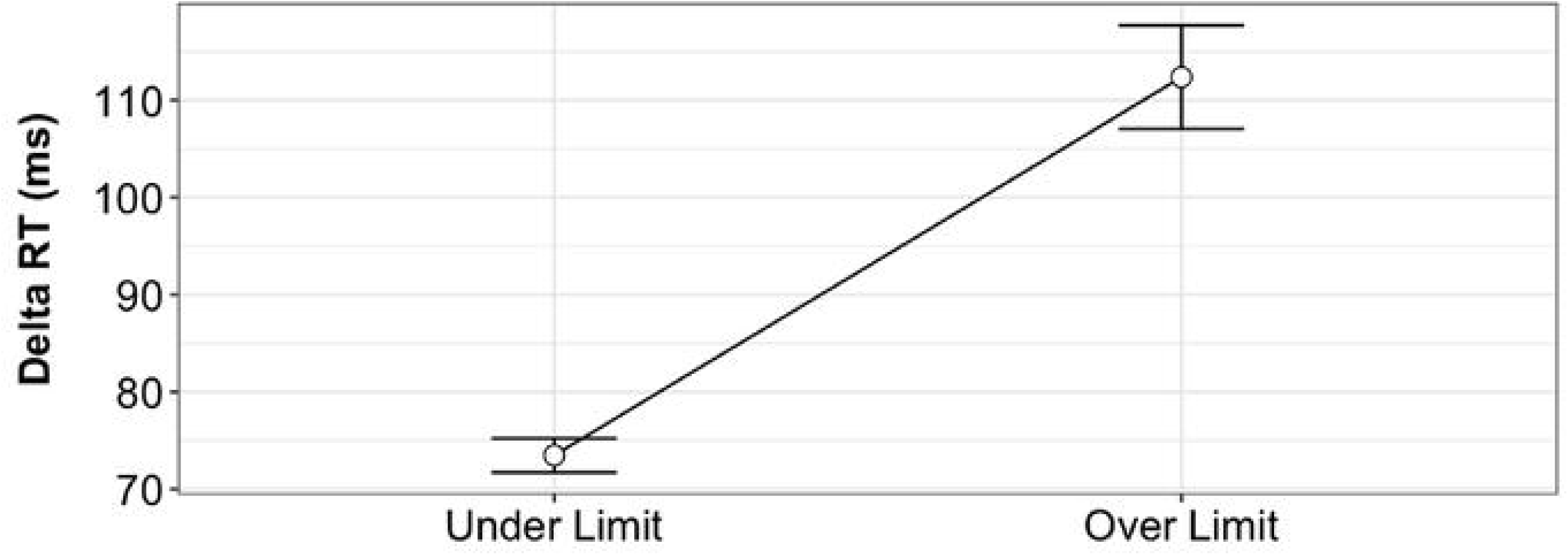
The separability of BAC level based on Delta RT. A one-way between-subjects ANOVA was conducted on the Delta RT for a group of participants while under the legal BAC limit (BAC <= 0.08%) and over the legal BAC limit (BAC > 0.08%). Pairwise comparisons show statistically significant differences between each factor at the p < 0.05 level.

A simple linear regression analysis was conducted to evaluate if BAC levels could predict the Delta RT between data from participants who were under or over the legal limit of 0.08%. Control participants are included in the analysis. A significant regression was found (*F*(1, 3547) = 323.3, *p* = 2.2e-16). The *R*^2^ was 0.0833, indicating that the BAC level explained approximately 8.3% of the variance in Delta RT. Figure 9 illustrates the results of this linear model.

**Figure 9:**
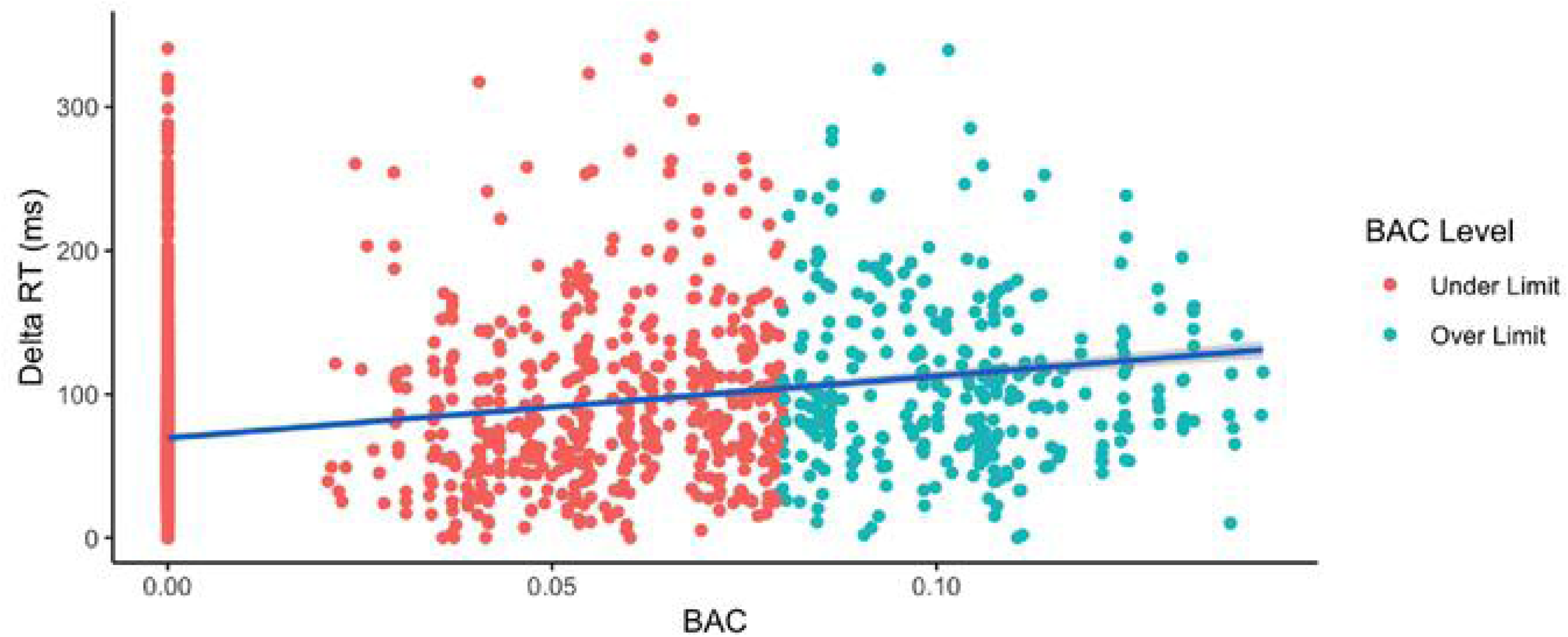
Linear regression analysis demonstrates the ability of blood alcohol content to predict changes in reaction time. Subject data was pooled for all participants, including controls, and is plotted in red for under the legal limit (BAC <= 0.08%) and in blue for over the legal limit (BAC > 0.08%) levels. A simple linear regression analysis was conducted to evaluate the extent to which BAC levels could predict the change in reaction time, denoted as “Delta RT.” A significant regression was found for the two conditions [F(1, 3547) = 323.3, p = 2.2e-16]). The *R*^2^ was 0.0833, indicating that the BAC level explained approximately 8.3% of the variance in Delta RT.

### Premotor Time and Blood Alcohol

A correlation of 0.874 (p<2.2e-16) was established between control subjects’ PMT and RT, shown in Figure 10, suggesting a consistent relationship between these two parameters and a consistent electromechanical delay. Changes in PMT were also assessed at an individual level over time as BAC increased. Mean PMT and BAC are plotted on separate axes as a function of time for two study participants and one control in Figure 11. Each data point in the PMT plot is the mean value across one three-minute measurement.

**Figure 10:**
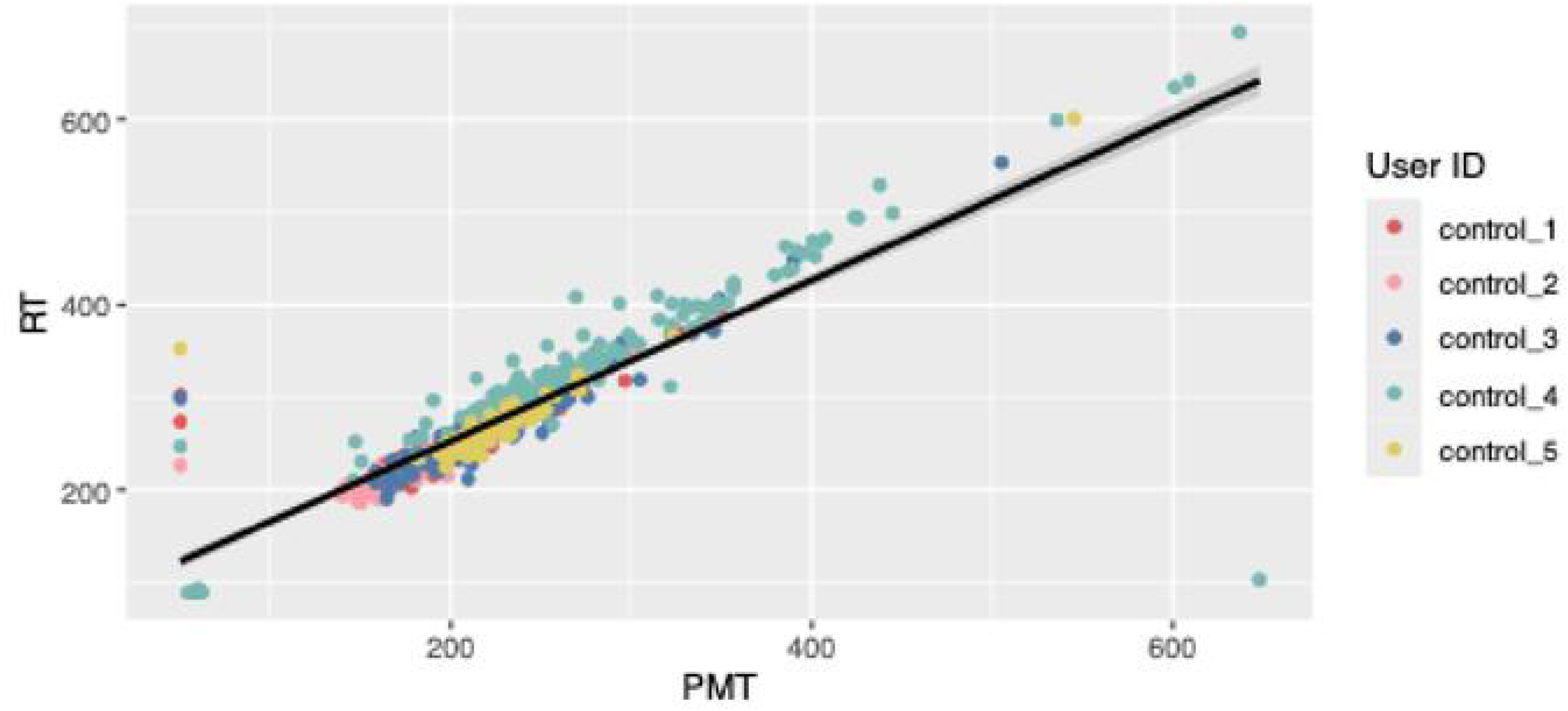
A correlation of 0.874 (p<2.2e-16) was established between premotor time and reaction time in control subjects, suggesting a consistent relationship and a consistent electromechanical delay.

**Figure 11.**
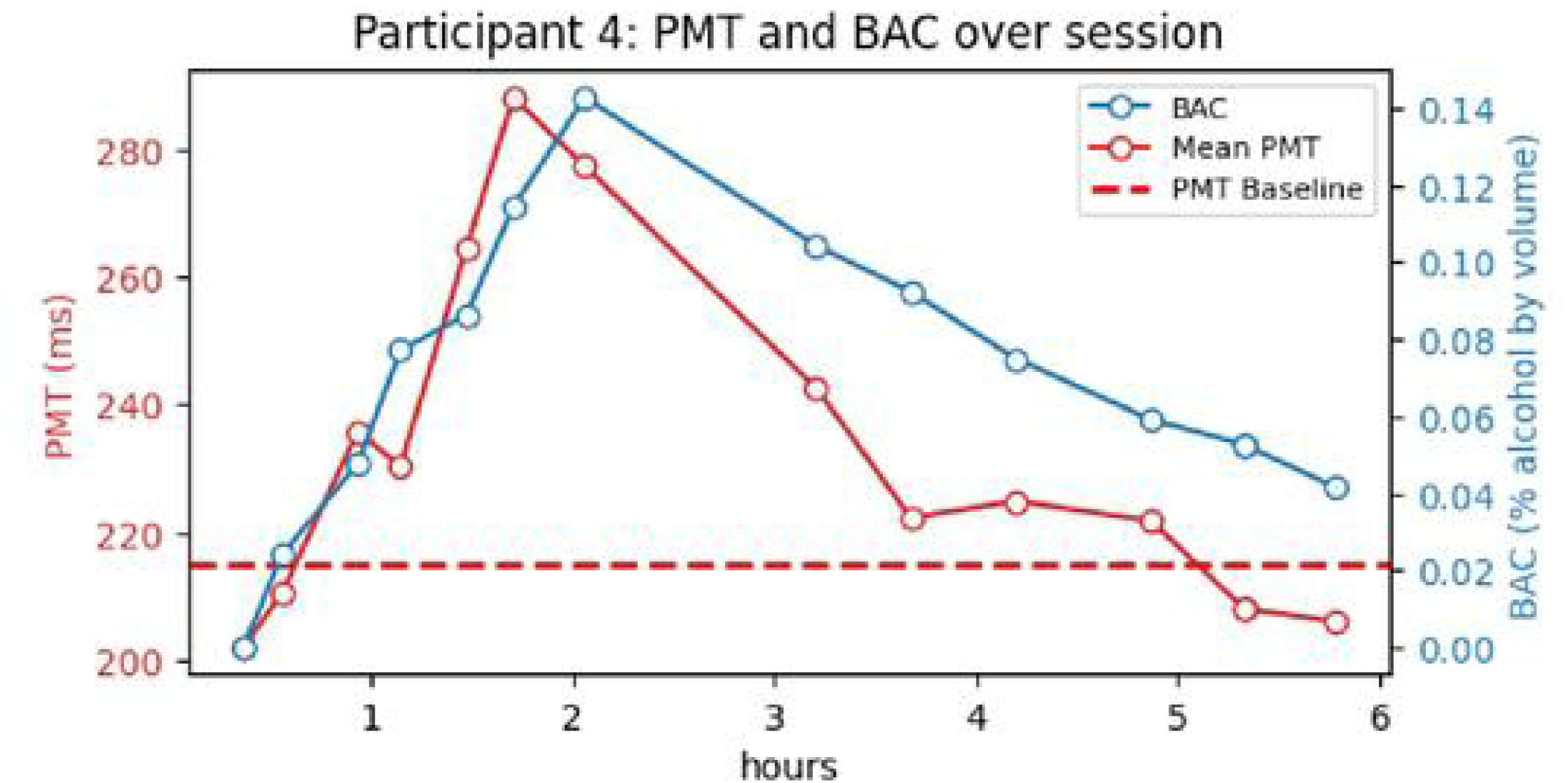
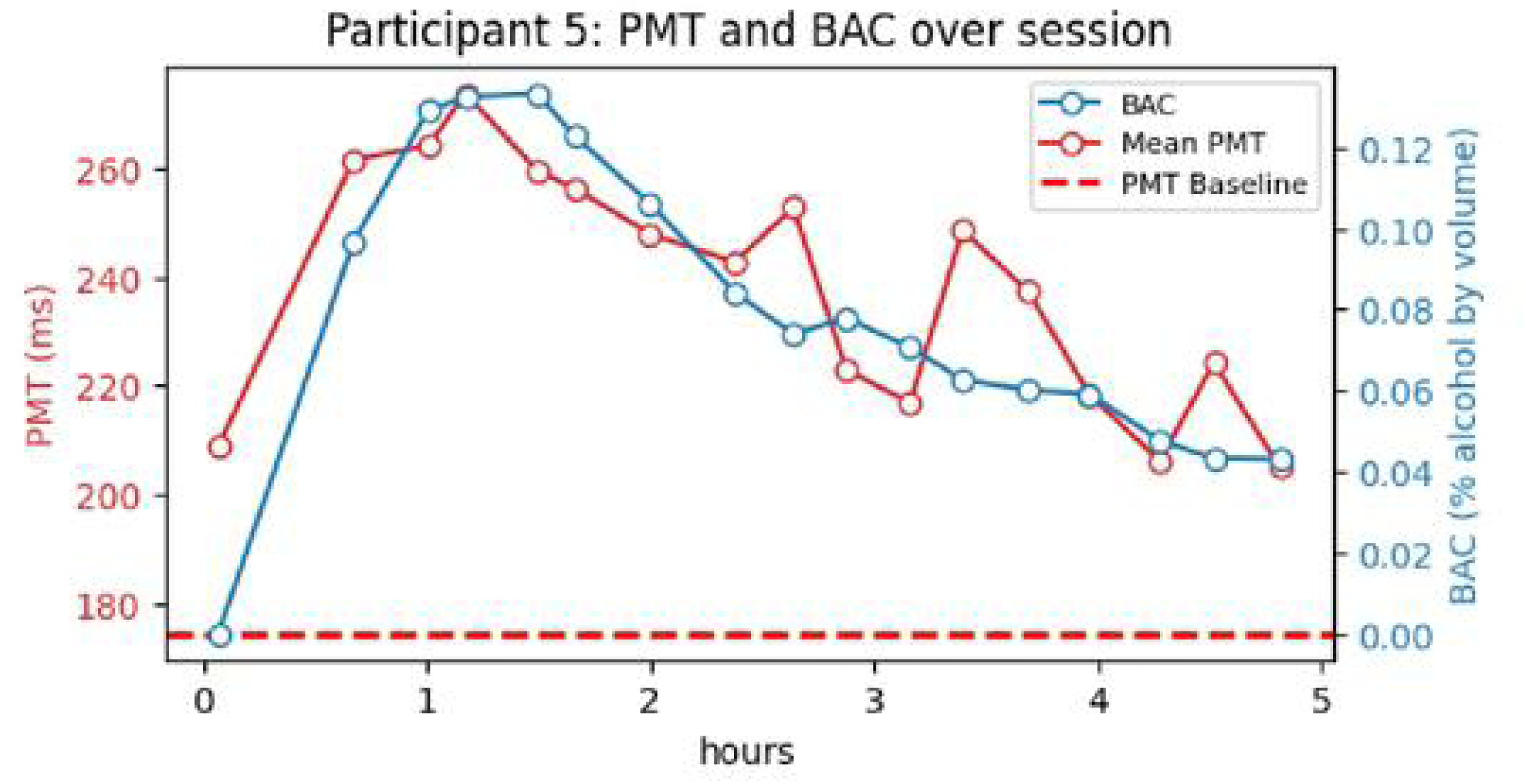
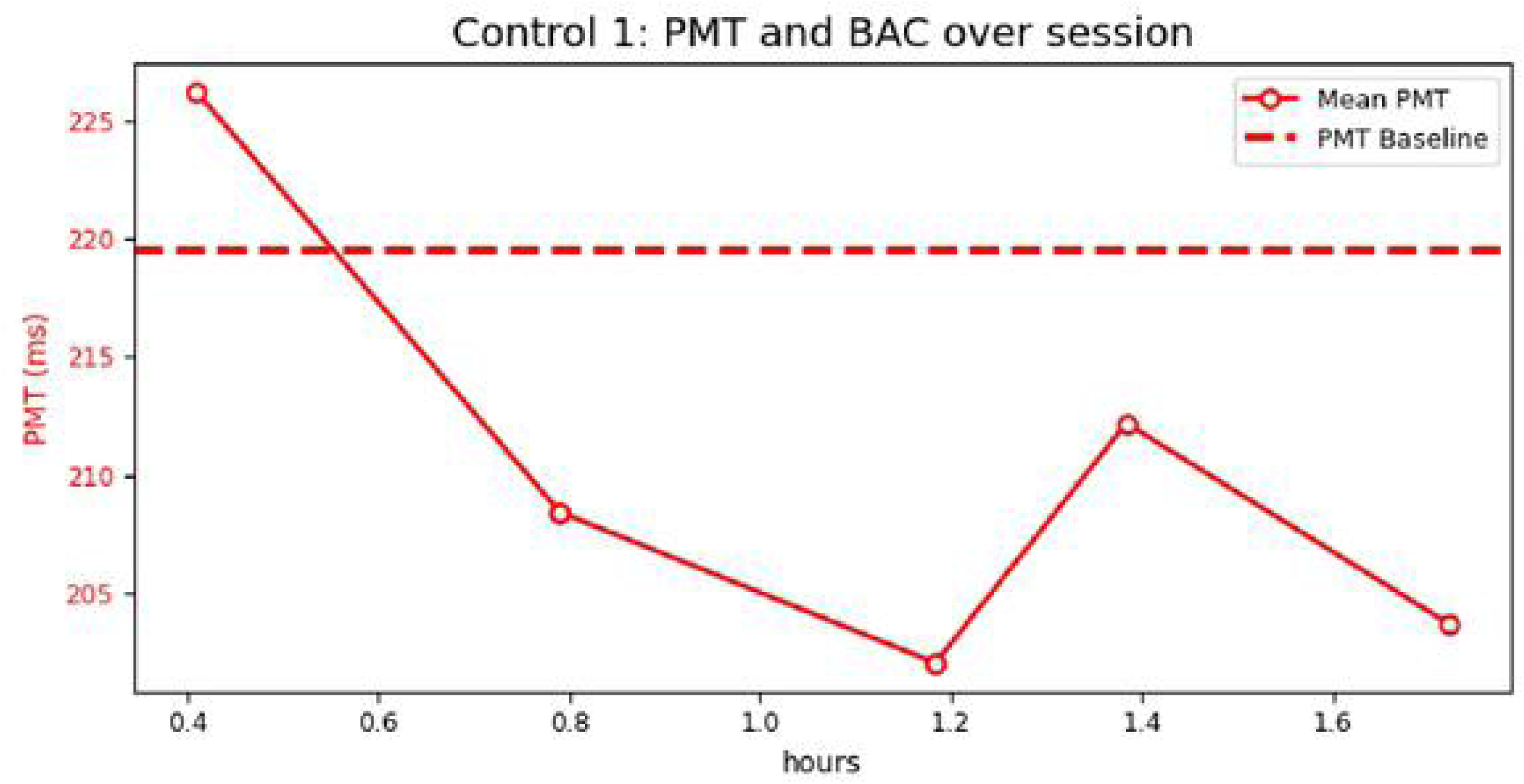
For each participant, blood alcohol and reaction time were measured before and intermittently after the alcohol dosage. In the top two plots (A and B), the mean PMT for each reaction time test was plotted over the session as well as the subjects’ BAC. The bottom plot (C) is for a control participant who did not consume alcohol. The mean PMT measured days before or days after the alcohol session was used to calculate the baseline PMT.

The baseline PMT is the average value of all results collected during an earlier session prior to the experimental condition and is denoted as a dashed red line on the plots. Data for all participants is shown in the Supplemental section. Most participants’ BAC and PMT peak between 1 and 2 hours post alcohol administration and display a similar rate of rise for the BAC and PMT values as well as a similar rate of return to baseline.

A repeated measures ANOVA was used to investigate changes in Delta RT over multiple measurements for participants who were not legally intoxicated (BAC<0.08), represented as the “Under Limit” level in Fig. 12, and for those same participants when they were legally intoxicated (BAC ≥ 0.08), represented as the “Over Limit” level in Fig. 12. The analysis showed a significant effect of BAC category on Delta RT, F(1, 2161) = 156, p = .0001, η² = 0.067, indicating that intoxication levels significantly impact reaction time changes. Seven of 13 participants (subjects 4, 5, 6, 7, 14, 17, and 22) demonstrated statistically significant differences in Delta RT between the two BAC conditions. These findings suggest that intoxication, as measured by BAC, significantly influences reaction time performance.

**Figure 12.**
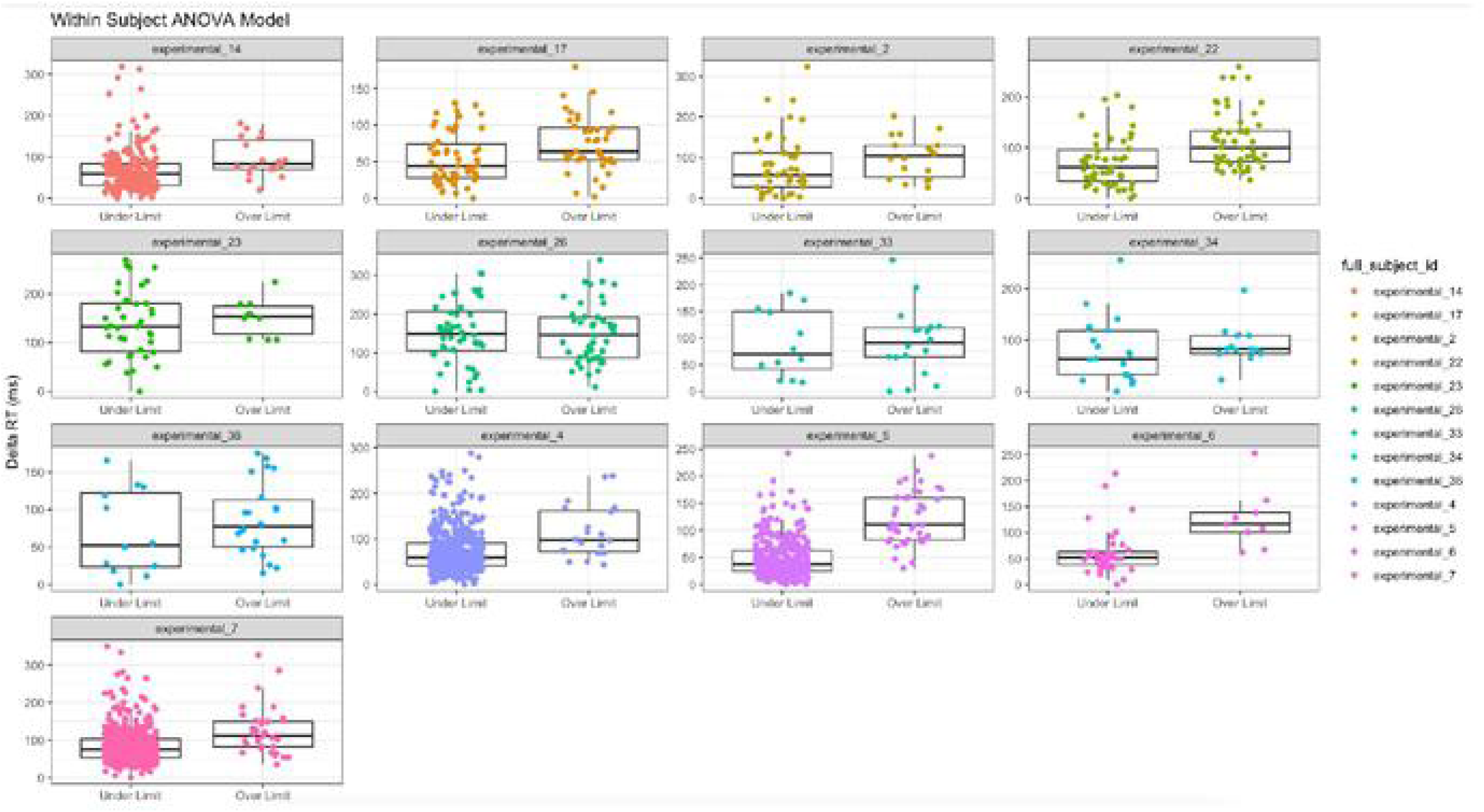
A repeated measures ANOVA showed a significant effect of BAC category on Delta RT. The “Under Limit” condition represents participants with BAC < 0.08 (not legally intoxicated), while the “Over Limit” condition represents the same participants when their BAC ≥ 0.08 (legally intoxicated). Seven of 13 participants (subjects 4, 5, 6, 7, 14, 17, and 22) demonstrated statistically significant differences in Delta RT between the two BAC conditions.

## Discussion

The Pison wearable device detected impairment in participants through measuring increases of reaction time concomitant with increases in blood alcohol concentration. Increases in premotor reaction time that occur after a moderate dose of alcohol are representative of a slowing of cognitive processes and have been demonstrated in prior laboratory-based studies using simultaneous EMG and EEG data collection.^19^ The pilot results presented in this study demonstrate a potential solution to reliably and accurately measure premotor time in real-world settings using a wearable having the footprint of a smartwatch. By monitoring reaction time unobtrusively in an easy-to-use device, individuals who are impaired by alcohol consumption can be prevented from operating a motor vehicle.

The neurophysiologic data measured with the Pison wearable provides a robust predictive capacity to determine, at a population level, in a statistically significant manner, the difference between subjects below and above the legal BAC. This has potential great significance in addressing the issue of impaired motor vehicle driving. Additionally, at an individual level, a very robust mirroring of reaction time to blood alcohol level was shown. This personalization of the data is extremely valuable in determining each individual’s cognitive ability to engage in critical tasks, such as driving.

Several limitations should be considered when interpreting these pilot study results. The sample size (n=19) was small and future studies should include larger numbers selected from heterogenous populations. Some participants collected baseline data in home settings, which may have differed in terms of ambient lighting or electromagnetic interference than the data collected in the laboratory setting. Future studies should have all participants collect data in real-world settings so that the impact of environmental factors can be assessed. Pison would like to conduct additional studies using its latest device, shown in Figure 13, that includes EMG and IMU neural sensors that are implemented on a readout integrated circuit developed in partnership with STMicro. Future studies will also compare the sensitivity of EMG and IMU measures to intoxication level as well as evaluate a composite measure of PMT and RT to identifying intoxication.

**Figure 13.**
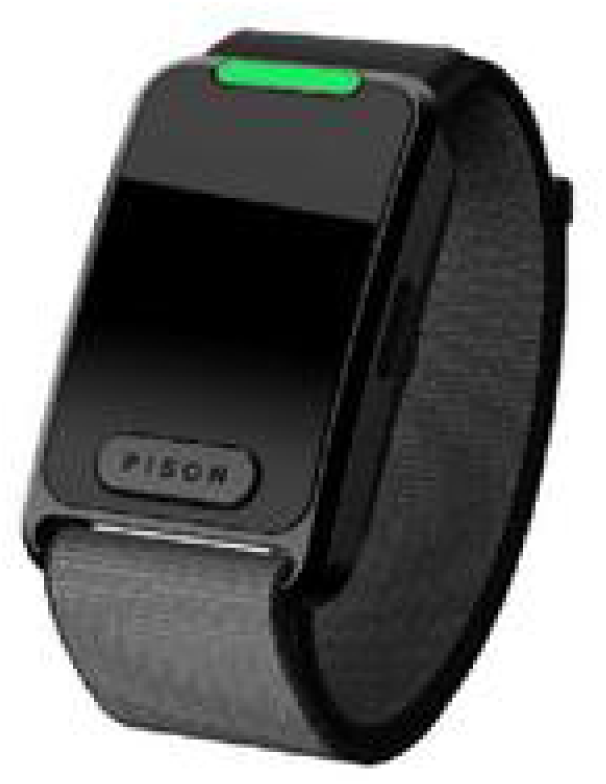
The Pison wearable sensor platform uses patented sensing technology to acquire biopotential, motion, and other physiological signals. Data from Pison’s surface electromyography, inertial measurement unit, photoplethysmography, electrocardiography, galvanic skin response, temperature, and ambient light sensors are transmitted to a client device (e.g., mobile phone) via Bluetooth Low Energy.

## Conclusion

In this study, the first on-body, real-time measurements of impairments associated with increasing blood alcohol content were conducted using neuro-physiologic Pison sensors. This study is the first to provide insight into the influence of neural sensors and computational power to inform impairment due to alcohol. Preventing motor vehicle use while under the influence of alcohol is an impactful use case and future studies will expand the subject size as well as broaden the sources of impairment beyond alcohol to include other legal substances such as tetrahydro cannabinol, illegal substance, as well as sleep deprivation or chronic fatigue. The use of reaction time can be used to measure impairment independent of the etiology of the cognitive decline.

These initial findings suggest a scalable solution to prevent driving under the influence of alcohol, exponentially reducing the risk to intoxicated drivers, passengers, other drivers, and pedestrians, thereby improving public safety. Additionally, the quantitative data produced by the Pison wearable may be used to inform casual drinkers of when they should not consume more alcohol to maintain sobriety.

## Acknowledgments

The authors would like to thank Camila Moreno-Bo and Valerie Novoa Chinchilla for their help formatting the manuscript. Cambridge Focus, Inc., and independent research participant recruiting agency was used for this study.

## References

1. Centers for Disease Control and Prevention. Alcohol and Public Health: Alcohol-Related Disease Impact (ARDI). U.S. Department of Health & Human Services, 2022. Available from: https://nccd.cdc.gov/DPH_ARDI/default/default.aspx

2. World Health Organization. *Alcohol Fact Sheet*, 2022. Available from: https://www.who.int/news-room/fact-sheets/detail/alcohol

3. Esser MB, Hedden SL, Kanny D, Brewer RD, Gfroerer BA, Naimi TS. Prevalence of Alcohol Dependence Among US Adult Drinkers, 2009–2011. Prev Chronic Dis. 2014; 11:140329. doi:10.5888/pcd11.140329.

4. National Center for Statistics and Analysis. Alcohol-impaired driving: 2020 data (Traffic Safety Facts. Report No. DOT HS 813 294), 2022. National Highway Traffic Safety Administration.

5. Weinberg A. Should Your Car Know When You’re Too Drunk to Drive? Mother Jones, 2023. Available from: https://www.motherjones.com/politics/2023/03/drunk-driving-infastructure-bill-technology-dadss-safety/

6. Pollard JK, Nadler ED, Stearns MD. Review of Technology to Prevent Alcohol-Impaired Crashes (TOPIC). U.S. Department of Transportation National Highway Traffic Safety Administration, 2007.

7. Guo X, Shojaei-Asanjan K, Zhang D, Sivajrurnathan K, Sun Q, Song P, et al. Highly Sensitive and Specific Noninvasive In-Vivo Alcohol Detection Using Wavelength-Modulated Differential Photothermal Radiometry. Biomed Opt Express. 2018; 9(10):4638–4648. doi:10.1364/BOE.9.004638.

8. Dinh TV, Choi IY, Son YS, Kim JC. A review on non-dispersive infrared gas sensors: Improvement of sensor detection limit and interference correction. Sens Act B: Chem. 2016; 231:529–538. doi:10.1016/j.snb.2016.03.040.

8. Bakowski DL, Savis ST, Moroney WF. Reaction time and glance behavior of visually distracted drivers to an imminent forward collision as a function of training, auditory warning, and gender. Procedia Manufacturing 2015; 3:3238–3245. doi:10/1016/j.promfg.2015.07.875.

9. Yadav AK, Velaga NR. Modelling the relationship between different Blood Alcohol Concentrations and reaction time of young and mature drivers. Transportation Research Part F 2019; 4:227–245. doi:10.1016/j.trf.2019.05.011.

10. McGehee DV, Mazzae EN, Baldwin GHS. Driver Reaction Time in Crash Avoidance Research: Validation of a Driving Simulator Study on a Test Track. Proc IEA 2000/HFES 2000 Congress. 2000; 44(20):3-320-3-323. doi:10.1177/154193120004402026.

11. Tzambazis K, Stough C. Alcohol impairs speed of information processing and simple and choice reaction time and differentially impairs higher-order cognitive abilities. Alcohol & Alcoholism. 2000; 35(2):197–201. 10.1093/alcalc/35.2.197.

12. Li YC, Sze NN, Wong SC, Yan W, Tsui KL, So FL. A simulation study of the effects of alcohol on driving performance in a Chinese population. Accident Analysis and Prevention. 2016; 95:334–342. doi:10.1016/j.aap.2016.01.010.

13. Liu YC, Ho CH. Effects of different blood alcohol concentrations and post-alcohol impairment on driving behavior and task performance. Traffic Injury Prevention. 2010; 11(4):334–341. doi:10.1080/15389581003747522.

14. Kuypers KPC, Samyn N, Ramaekers JG. MDMA and alcohol effects, combined and alone, on objective and subjective measures of actual driving performance and psychomotor function. Psychopharmacology. 2006;187(4):467–475. doi:10.1007/s00213-006-0434-z.

15. Norman RW, Komi PV. Electromechanical delay in skeletal muscle under normal movement conditions. Acta Physiol Scand. 1979;106(3): 241–248. doi: 10.1111/j.1748-1716.1979.tb06394.x.

16. Holden J, Francisco E, Tommerdahl A, Lensch R, Kirsch B, Zai L, et al. Methodological problems with online concussion testing. Frontiers in Human Neuroscience. 2020;14, Article 509091. doi:10.3389/fnhum.2020.509091.

17. Estimated BAC Charts. BRAD Foundation. [Cited 15 September 2023]. Available from: https://brad21.org/estimated-bac-charts

18. Hernandez OH, Vogel-Sprott M. Alcohol slows the brain potential associated with cognitive reaction time to an omitted stimulus. J Stud Alcohol Drugs. 2010;71(2):268–77.

